# Dysregulated *MEG3* in Myotonic Dystrophy 1: nuclear retention, pathological role, and therapeutic correction by antisense conjugates

**DOI:** 10.1101/2025.06.20.658327

**Authors:** David Seoane-Miraz, Jessica Stoodley, Natalia Galindo Riera, Yahya Jad, Arnaud F. Klein, Lara Nikel, Yulia Lomonosova, Jesus Reine, Susana Camara, Ruben Artero, Denis Furling, Matthew JA Wood, Miguel A Varela

## Abstract

*MEG3*, a long non-coding RNA (lncRNA), has been shown to play a critical role in regulating apoptosis. Its downregulation inhibits apoptosis in cancer cells, whereas its upregulation has been associated with cell death in both cardiovascular disease and, more recently, Alzheimer’s Disease. Here we show that *MEG3* is upregulated in Myotonic Dystrophy 1 (DM1). Specifically, we show MEG3 upregulation by several-fold in DM1 human muscle cells and in two DM1 mouse models, HSA-LR and LC15. In human DM1 muscle cells we observe nuclear retention of MEG3 and an increase in its transcript diversity. Furthermore, we observe a general trend of nuclear retention in DM1 affecting lncRNAs and microRNAs (miRNAs), in contrast to mRNAs, when compared to healthy cells. This altered nuclear retention may contribute to the pathological effects of non-coding RNA dysregulation in DM1. Importantly, we demonstrate that treatment with antisense conjugates targeting the repeat expansion causing DM1, an approach currently being tested in Clinical Trials, corrects *MEG3* levels in HSA-LR mice, without additional therapeutic interventions targeting *MEG3*.

## Introduction

Myotonic dystrophy type 1 (DM1) is a neuromuscular disorder caused by a repeat expansion in the *DMPK* gene. This expansion causes the sequestration of splicing factors such as muscleblind-like splicing regulator 1 (*MBNL1*) by mutant expanded RNA which form foci in the nucleus, leading to genome-wide mis-splicing events (1). DM1 symptoms are characterised by progressive muscle weakness and myotonia. However, the molecular causes behind this loss of muscle cells in DM1 and other neuromuscular diseases are complex and poorly understood.

In this study, we focus on the lncRNA *MEG3* (maternally expressed gene 3). This transcript promotes cell apoptosis and has been studied in the context of cancer (2). Furthermore, *MEG3* has been linked to neuronal loss in Alzheimer’s Disease (3) and the promotion of apoptosis in cardiovascular disease models (4, 5).

Our data reveal a striking retention of *MEG3* in DM1 nuclei, suggesting a fundamental defect in lncRNA and miRNA transport in DM1 pathology, in contrast to mRNAs, for which the opposite trend is detected. These changes in transcript localization could exacerbate the pathological effects of some non-coding RNAs and be at least partially responsible for the upregulation of many genes that code for mRNA previously described in DM1 (6). Importantly, it demonstrates that treatment with antisense conjugates, targeting the repeat expansion causing DM1 can correct elevated levels of *MEG3* without the need of additional therapeutic compounds.

## Results and Discussion

### lncRNAs and miRNAs are retained in the nucleus of human DM1 muscle cells contrary to mRNA

RNA-Seq data produced from the nucleus and cytoplasm of DM1 human muscle cells containing CUG expanded repeats in the *DMPK* gene (2,600 repeats) showed an enrichment of lncRNAs and miRNAs retained in the nuclear fraction of DM1 cells in comparison to wild-type counterparts, a phenomenon not observed in mRNAs (Fig. 1). These data suggests that nuclear retention of lncRNA and miRNAs could be playing a role in DM1 pathogenesis. We found no evidence that the retained transcripts are sequestered in the nucleus by the mutant *DMPK* nuclear foci. Similarly, we saw no sequence enrichment in the differentially retained miRNAs or lncRNAs in the nuclear compartment of DM1 muscle cells (see extended Methods in Supporting Information). However, extensive mis-splicing events existed in DM1 cells, including lncRNAs and genes involved in RNA trafficking and stability, and *MEG3* transcript diversity and nuclear localization was greatly increased (Fig. 1B). We confirmed *MEG3* upregulation by qPCR, revealing a 200-fold increase in the cytoplasm and a >2000-fold increase in the nucleus of DM1 human muscle cells when compared to wild-type counterparts (Fig. 2).

**Figure 1.**
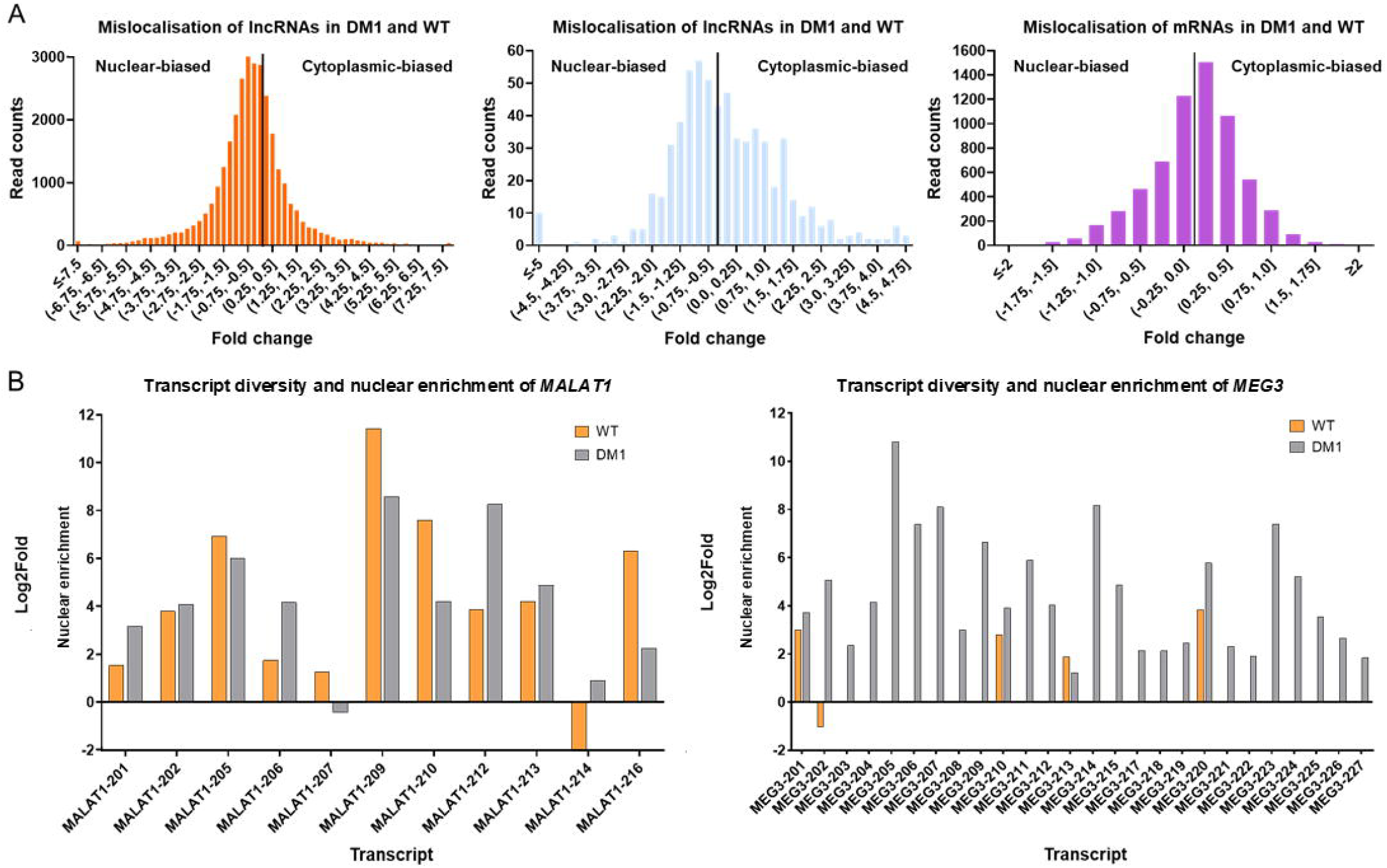
The transcriptome of DM1 patient-derived muscle cells shows nuclear retention of non-coding RNA in contrast to mRNA and upregulation and missplicing of *MEG3*. Transcriptomic analysis by RNA sequencing was performed on total RNA isolated from DM1 myotubes compared to wild type (n = 3). (A) Nuclear accumulation of lncRNAs and miRNAs in DM1 muscle cells compared to wild-type cells. Normalized read counts revealed a global shift in the abundance of transcripts between the nucleus and cytoplasm in DM1. In DM1, there was a 36.4% increase in lncRNAs upregulated in the nucleus and a 28.3% increase in miRNAs (p-value adjusted for multiple comparisons, padj ≤ 0.05; log2 Fold Change ≥ 0). This accumulation in the nucleus of DM1 cells was not observed for mRNA. (B) Increased diversity in *MEG3* transcripts and nuclear localization.

**Figure 2.**
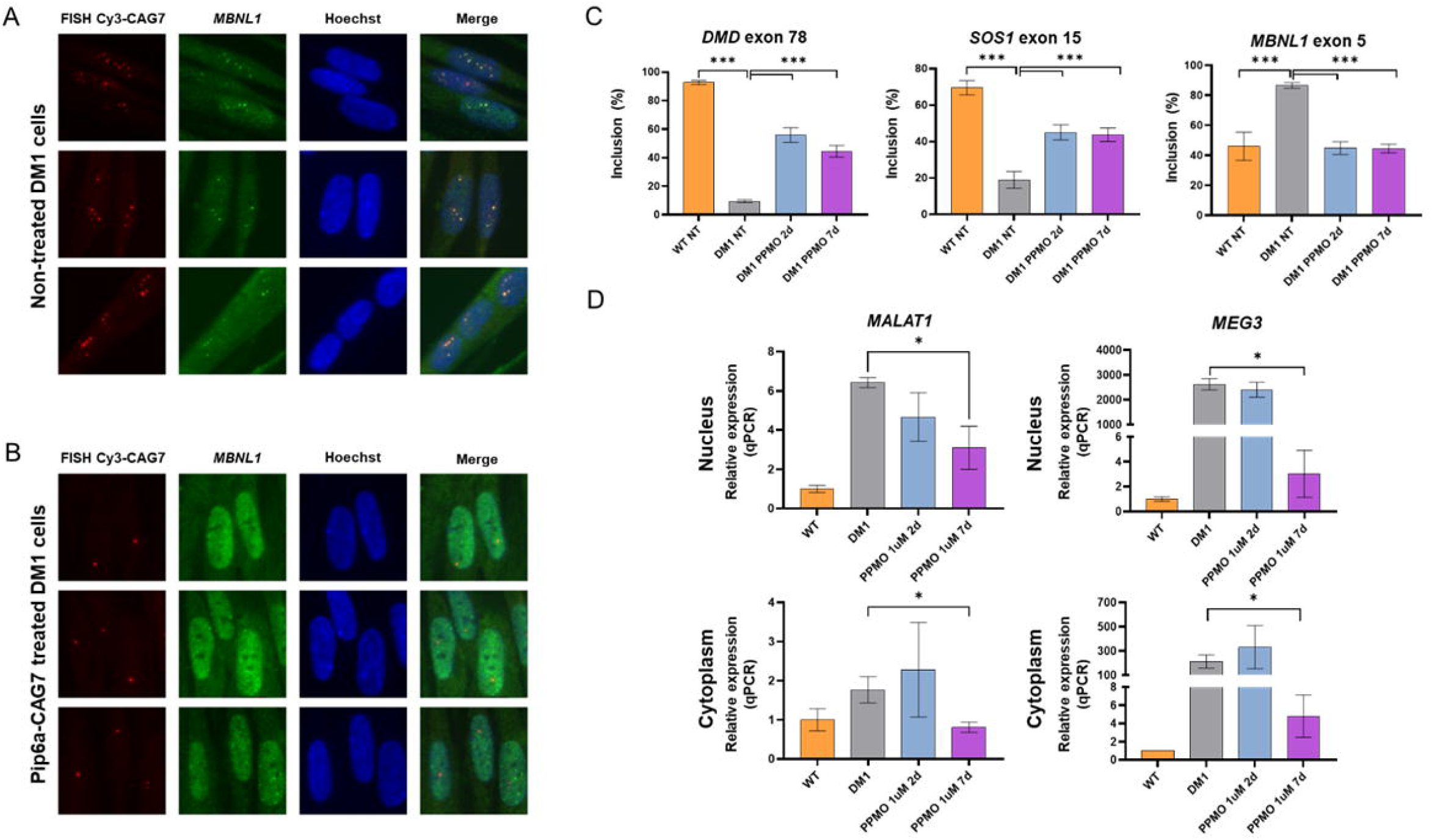
Antisense conjugates correct *MEG3* upregulation in DM1 patient-derived muscle cells. Human DM1 myotubes with a 2600 CTG repeat expansion were treated with 1 μM of the peptide-PMO conjugate Pip6a-CAG7 targeting the repeat expansion and analysed 24 h later. (A-B) FISH shows a reduction in the number of toxic RNA foci after treatment, leading to the resulting desequestration of the splicing factor MBNL1, (C) RT-PCR analysis of mis-splicing events and correction after 2 or 7 days after 1 µM Pip6a-CAG7 treatment. (D) *MALAT1* and *MEG3* correction was assessed by RT-qPCR 2 or 7 days after treatment at 1 µM. Data are expressed as mean ± SEM. * P < 0.05, *** P < 0.001 by 1-way ANOVA with Newman-Keuls post hoc test.

In healthy cells, the presence of motifs recruiting RNA binding proteins, inefficient splicing or RNA instability can lead to poor nuclear export (7). *MEG3* contains a nuclear retention element (NRE) for promotion of MEG3 localisation within the nucleus (8). In DM1 cells, expression, transcript diversity, and nuclear retention of *MEG3* transcripts are increased potentially exacerbating *MEG3* function. Zuckerman and Ulitsky’s (9) analysis of nuclear-cytoplasmic RNA localization showed splicing efficiency as the predominant predictive factor in both wild-type mice and human cells. In DM1, wide mis-splicing and nuclear retention give support to a two-step nuclear retention model where the initial accumulation of expanded *DMPK* RNA in nuclear foci sequester proteins involved in splicing deregulating both splicing and RNA transport efficiency, leading to widespread nuclear retention of lncRNAs and miRNAs.

### MEG3 upregulation correlates with muscle pathology and is rescued by antisense conjugates targeting the transcripts

The significant increase in DM1 of *MEG3* expression prompted the consideration of this lncRNA as a potential therapeutic target for DM1. We measured Meg3 by qPCR in two DM1 mouse models, HSA-LR and LC15, in comparison with their background wild-type (FVB/N). Figure 3 shows that Meg3 is upregulated in the skeletal muscle of both models, with a 15-fold upregulation of Meg3 in HSA-LR and a 6-fold upregulation in LC15. Together with the upregulation of MEG3 in human cells from our *in vitro* data, that demonstrates a correlation of *MEG3* levels with muscle disease, this result led us to hypothesise that *MEG3* correlates with muscle disease in DM1.

**Figure 3.**
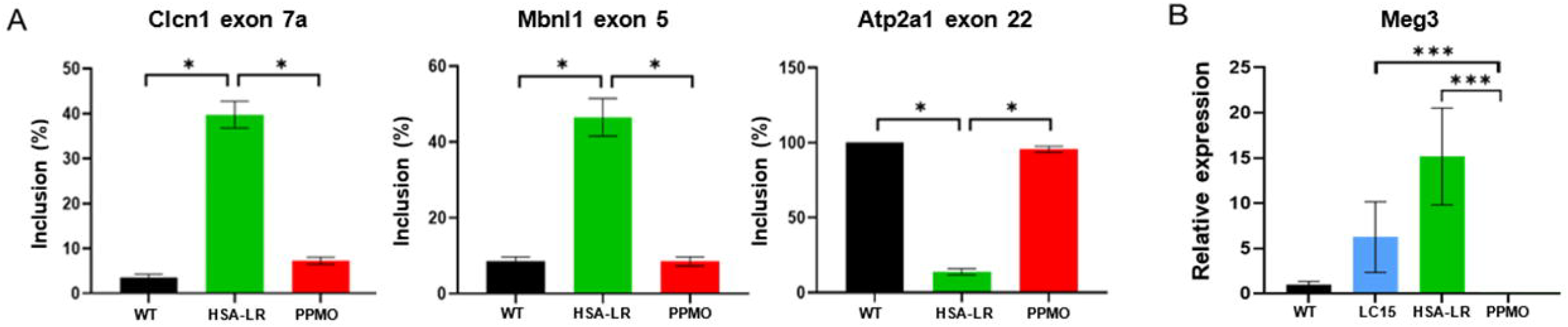
Meg3 is upregulated in two DM1 mouse models and is corrected by antisense conjugates targeting RNA *foci*. HSA-LR and LC15 mice were injected in the tail vein with three 12.5 mg/kg doses of the peptide-PMO conjugate Pip6a-CAG7 targeting the repeat expansion administered intravenously twice per week and analysed two weeks post-treatment (n = 4). (A) RT-PCR analysis of mis-splicing in quadriceps for DM1 biomarkers Clcn1, Mbnl1 and Atp2a1 shows the correction of splicing events to wild-type levels 2 weeks after three doses of Pip6a-CAG7. (B) Meg3 is upregulated by 15-fold in the quadriceps of HSA-LR and 6-fold in LC15, relative to wild-type controls (FVB/N), as measured by qPCR normalised to the average expression of Gapdh and Actb. Treatment with antisense conjugates targeting the repeat expansion causing DM1 corrects Meg3 levels without additional therapeutic interventions targeting Meg3. Data are expressed as mean ± SEM. * P < 0.05, *** P < 0.001 by 1-way ANOVA with Newman-Keuls post hoc test.

To test our hypothesis, we employed peptide-antisense conjugates known to correct DM1 pathology in patient-derived muscle cells and the mis-splicing and myotonia phenotype of HSA-LR mice (6). Peptide-antisense conjugates targeting the repeat sequence work through a steric blocking mechanism, preventing the sequestration of the splicing factor *MBNL1*. In human DM1 myotubes with a 2,600 CTG repeat expansion, FISH analysis revealed a reduction in the number of toxic RNA foci following treatment, which led to the desequestration of *MBNL1* after treatment with 1 µM of the antisense conjugate Pip6a-CAG7 (Fig. 2A). Additionally, significant corrections were observed in DM1 splicing biomarkers, including *DMD, SOS1*, and *MBNL1*. Notably, these conjugates also restored MEG3 expression in both the nucleus and cytoplasm seven days post-treatment (Fig. 2B).

To assess whether a similar correction of Meg3 could be achieved *in vivo*, we administered three biweekly intravenous doses of 12.5 mg/kg Pip6a-CAG7 in the DM1 model HSA-LR (6). Two weeks after the last administration we observed the complete correction of Meg3 levels. We next evaluated the possibility that Meg3 remained upregulated after a suboptimal dose (7.5 mg/kg), producing only 50% myotonia correction two weeks post-administration. Again, Meg3 levels decreased to wild-type levels (Fig. 3). These results show that Meg3 correlates with muscle disease in DM1 and that antisense treatments targeting the mutated Dmpk transcript also robustly downregulate Meg3 without additional therapeutic interventions.

## Materials and Methods

Peptide-oligonucleotide conjugates were synthesised as described in (10). All animal procedures were authorised by the UK Home Office and by the University of Oxford ethics committee (PPL no: PDFEDC6F0). For detailed methods, please see SI Appendix.

## Author Contributions

D.S.M. and M.A.V. conceived the study. D.S.M., J.S., N.G.R., Y.J., A.F.K. and L.N. performed experiments and acquired data. D.S.M., L.N., N.G.R., Y.L. and M.A.V., analyzed the data. R.A., D.F., M.J.A.W and M.A.V. supervised the study. D.S.M. and M.V. wrote the first draft of the manuscript. All authors contributed to the final draft of the manuscript.

## Acknowledgments

This work was supported by Medical Research Council grants (MRW0147421 and MRP01741×1), John Fell Fund (0007895), Muscular Dystrophy UK (19GRO-PG36-0294), Margarita Salas Fellowship awarded to NG and Association Francaise contre les Myopathies/AFM-Telethon (grants 15758 and 22922).

