## Supplementary Information for "Dysregulated *MEG3* in Myotonic Dystrophy 1: nuclear retention, pathological role, and therapeutic correction by antisense conjugates"

Miguel A. Varela

Supporting Information Methods

*Synthesis of peptide-phosphorodiamidate morpholino oligomer (P-PMO) conjugates*

Penetrating peptide Pip6a (Ac-(RXRRBRRXRYQFLIRXRBRXRB)-COOH) was synthesised by standard Fmoc solid-phase chemistry and covalently attached to amine in 3ʹ terminus of the phosphorodiamidate morpholino oligomer (PMO) targeting CTG expanded repeats (5′-CAGCAGCAGCAGCAGCAGCAG-3′) (Gene Tools LLC) as described in Stoodley et al. (1).

*Human-derived myotube cultures*

Immortalised primary muscle cells derived from healthy (wild-type control) and DM1 patients carrying a mutation with 2,600 CTG repeats were obtained from Arandel et al. (2). Myoblasts were cultivated in Skeletal Muscle Cell Growth Medium (PromoCell) supplemented with 1% penicillin/ streptomycin/ amphotericin B in an incubator at 37 ºC and an atmosphere of 5% CO_2_, until cell confluence reached approximately 80%, changing the media on alternate days. Myoblast differentiation into myotubes was then induced by replacing growth media with Skeletal Muscle Cell Differentiation Medium (PromoCell) for 5 to 8 days, changing the media on alternate days.

*Mouse models and P-PMO injections*

In this study we use two DM1 mouse models: HSA-LR and LC15. HSA-LR is the most used mouse in DM1 preclinical development. It displays myotonia and robust splicing changes from the expression of a transgene (human *ACTA1*) specifically in skeletal muscle. The transgene contains an insertion of ~250 CUG repeats in the 3’ untranslated region (3). LC15 mice ubiquitously express a human *DMPK* transgene and display milder splicing changes (4). Both mouse models have FVB/N wild type background. Pip6a administration in HSA-LR mice was performed by single intravenous injections in the tail vein at a dose of 7.5 mg/kg. Animals were culled 14 days after injection by cervical dislocation and tissues were harvested and kept at –80 ºC until analysis. All animal work was conducted according to procedures authorized by the UK Home Office under the Animal (Scientific Procedures) Act 1986.

*Cell fractionation, miRNA and lncRNA isolations*

Wild-type control and DM1 myotubes were harvested for RNA isolation once differentiation was completed. For miRNAs isolation, wild-type control and DM1 myotubes were fractionated into nucleus and cytoplasm using PARIS kit (Invitrogen) until the step in which both sub-cellular fractions were separated, followed by subsequent miRNAs isolation using miRNeasy Mini Kit (Qiagen) following manufacturer’s instructions. For lncRNAs isolation, similarly to miRNAs fractionation, wild-type control and DM1 myotubes were separated into nucleus and cytoplasm fractions using PARIS kit (Invitrogen), and total RNA was isolated by RNeasy Mini Kit (Qiagen) following manufacturer’s instructions. RNA quality and concentration was determined using a NanoDrop 2000 (Thermo Scientific) spectrophotometer.

*Library preparation and RNA sequencing*

Total RNA and miRNA enriched RNA from wild-type control and DM1 myotubes were sent to the Oxford Genomics Centre for sequencing. For miRNA sequencing, cDNA libraries were prepared using a TruSeq Small RNA Library Preparation Kit (Illumina) following manufacturer’s instructions. Sequencing was performed using rapid mode in a HiSeq 2500 System (Illumina). For lncRNA sequencing, cDNA libraries were prepared using an Illumina Stranded Total RNA Prep with Ribo-Zero Plus (Illumina), according to manufacturer recommendations. The total RNA was ribodepleted and this fraction was converted to cDNA. A second strand cDNA synthesis incorporated dUTP before the cDNA was end-repaired, A-tailed and adapter-ligated. Prior to amplification, samples underwent uridine digestion and prepared libraries were size selected, multiplexed and quality checked before sequencing. Paired-end sequencing was performed in a HiSeq 4000 System (Illumina).

RNA-Seq bioinformatic analysis

Single-end reads from miRNA library were cleaned from adaptors using Trimmomatic (5) and quality checked with FastQC application to remove from the dataset those reads with low quality values (Phred score threshold below 20). Follow-up analyses were computed through webserver sRNAtoolbox (6) including reads assembling, reads mapping on human GRCh38_p12 reference genome, reads annotation based on miRBase reference database, reads count, and differential expression analysis were carried out in DESeq2 (7). Computational analyses for paired-end reads from lncRNA libraries included quality control and filtering reads assembling and mapping to human reference genome. LncRNA identification and quantification, and differential expression were analysed using DESeq2 (7). We used Tandem Repeat Finder (8) to search for repetitive sequence enrichment whether uninterrupted or forming sequences with cryptic simplicity in the differentially retained miRNAs or lncRNAs in nuclear DM1 muscle cells. Complete raw data generated from RNA sequencing were deposited in the NCBI’s Gene Expression Omnibus database, GSE262163 (lncRNa and mRNA) and GSE262164 (miRNA).

*Validation of miRNA expression levels by RT-qPCR*

Preparation of 2 ng of cDNA templates was carried out using TaqMan Advanced miRNA cDNA Synthesis Kit (Applied Biosystems) following manufacturer’s instructions. This cDNA template was used subsequentially for quantification of mature miRNAs in triplicate in a StepOnePlus Real-time PCR system (Applied Biosystems). TaqMan assays (ThermoFisher Scientific) used in this study included the following: miR-1 (ID477820_mir), miR-133a-3p (ID478511_mir), miR-133a-5p (ID478706_mir), miR-133b (ID480871_mir), miR-206 (ID477968_mir), miR-493-3p (IDID478943_mir), and miR-504-5p (ID478144_mir) as targets, and miR-26b-5p (ID478418_mir), and miR-423-5p (ID478090_mir) as controls. qPCR reaction consisted of 1 cycle of 20 s at 95ºC for polymerase activation and 40 cycles of 1 s at 95ºC for cDNA denaturalisation and 20 s at 60ºC for annealing and extension.

*Validation of lncRNA and mRNA expression levels by RT-qPCR*

Reverse transcription of 2 ng of total RNA was performed using High-Capacity cDNA Reverse Transcription Kit (Applied Biosystems), following manufacturer’s instructions. Then, 2 ng of cDNA were used to quantify lncRNA and mRNA expression levels by qPCR using TaqMan Fast Advanced Master Mix for qPCR (Applied Biosystems). Following manufacturer’s recommendations, each reaction contains 5.0 µL of TaqMan Fast Advanced Master Mix (2X), 0.5 µL of TaqMan Assay (20X), 3.5 µL of nuclease-free water and 1 µL (2 ng) of cDNA template. qPCRs were carried out in triplicate in a StepOnePlus Real-time PCR system (Applied Biosystems) consisting of 1 cycle of 20 s at 95ºC for polymerase activation and 40 cycles of 1 s at 95ºC for cDNA denaturalisation and 20 s at 60ºC for annealing and extension.

*Fluorescent in situ hybridization (FISH) and immunofluorescence assay*

Combined FISH and immunofluorescence experiments were performed as described in Stoodley et al. (1) using a Cy3-labelled 2′OMe CAG7 probe (IDT) and a primary mouse monoclonal anti-*MBNL1* antibody (Developmental Studies Hybridoma Bank, Antibodies at the University of Iowa for use in research) followed by a secondary AlexaFluor 488 goat anti-mouse IgG, IgM (H+L) antibody (ThermoFischer Scientific). Images were taken using an Olympus BX60 microscope and Metamorph software (Molecular Devices). Confocal images were captured with a Nikon Ti2 microscope equipped with a motorized stage and a Yokogawa CSU-W1-spinning disk head coupled with a Prime 95 sCMOS camera (Photometrics). Images were processed with Adobe Photoshop software.

**Supporting Information References**
